# Meta-analysis shows that context-dependent mating behaviour is inconsistent or weak across animals

**DOI:** 10.1101/2020.08.23.263244

**Authors:** Liam R. Dougherty

## Abstract

Animals often need to invest significantly in mating behaviour in order to successfully mate. However, the expression of mating behaviour can be costly, especially in unfavourable environments, so animals are expected to adjust their behaviour in a context-dependent way to mitigate these costs. I systematically searched the literature for studies measuring animal mating behaviour (sexual signalling, response to sexual signals, or the strength of mate choice) in more than one environment, and used a phylogenetically-controlled meta-analysis to identify environmental factors influencing these behaviours. Across 222 studies, the strength of mate choice was significantly context-dependent, and most influenced by population density, population sex ratio, and predation risk. However, the average effect sizes were typically small. The amount of sexual signalling and the strength of response to sexual signals were not significantly related to the environment. Overall, this suggests that the evidence for context-dependent mating behaviour across animals is surprisingly weak.

## Introduction

For sexual animals, reproduction requires successfully mating with an individual of the opposite sex. In order to achieve this, individuals may need to signal or display to potential partners in order to attract and court them, or respond to the signals or displays of others. Additionally, some individuals make better mates than others, so that animals may gain considerable benefits from choosing only to mate with partners of the highest quality, leading to the expression of mate choice (Andersson 1994; Kokko *et al*. 2003; Rosenthal, 2017). However, both sexual signalling, and responding to such signals, can be expensive in terms of time and energy (Andersson 1994; Kotiaho 2001). There are also costs associated with mate choice, such as the energy and time needed to sample mates effectively (Sullivan 1994; Vitousek *et al*. 2007), or the risk of failing to mate if individuals are overly choosy (Barry & Kokko 2010; Greenway *et al*. 2015). In general, we expect the expression of these mating behaviours to be influenced by the balance of these costs and benefits: a behaviour should only be expressed when the benefits outweigh the costs.

Importantly, the costs and benefits of investing in these mating behaviours are inherently linked to the social, biological or physical environment. For example, at high predator density the cost of mate searching or sexual signalling is increased, if such behaviours make signallers or searchers more conspicuous (Magnhagen 1991; Zuk & Kolluru 1998). In these conditions, animals may benefit from investing less into searching and signalling, at least in the short-term. Importantly, the natural environment is complex, fluctuating, and unpredictable, both spatially and temporally (Miller & Svensson 2014). Therefore, animals will maximise their fitness by identifying situations in which mate searching and choice are beneficial or costly, and changing their behaviour accordingly. Indeed, evidence from a wide range of species shows that individuals often alter their mating behaviour over the short-term, in response to a wide range of social, biological, or physical factors (Jennions & Petrie 1997; Ah-King & Gowaty 2016; Kelly 2018). For example, many species respond to an increased predation risk by reducing signalling (e.g. Endler 1987; Fuller & Berglund 1996) or exhibiting weaker mate choice (e.g. Hedrick & Dill 1993; Gong & Gibson 1996; Hughes et al. 2012).

These empirical examples show that environment can be an important determinant of mating behaviour in some species. Importantly, by identifying these effects in laboratory studies, we may be able to better predict the expression of mating behaviour in the natural environment, which is complex and highly dynamic (Miller & Svensson 2014). Further, mate choice is a key component of sexual selection, which can influence population fitness and drive the evolution of novel phenotypes, the action of which may in turn be influenced by the expression of sexual signals (Andersson 1994). Therefore, understanding the extent to which both signalling and mate choice are context-dependent will help us to predict the strength of sexual selection, and the resulting evolutionary change, in natural populations. However, such predictions will only be possible if environmental effects are generally consistent across species, and there is evidence that this may not be the case. For example, many studies fail to find any significant effect of the environment on mating behaviour (e.g. in relation to predation risk: Briggs et al. 1996; Billing et al. 2007). In other cases, studies find significant effects, but in contrasting directions (e.g. Beckers & Wagner 2018), further suggesting that environmental effects on mating behaviour may not be as clear as previously suggested. Importantly, to date there has been no quantitative synthesis of these data.

To address this problem, I systematically searched for studies reporting animal mating behaviour in relation to seven environmental factors that are predicted to influence the costs and benefits of expressing these behaviours. In order to estimate the degree of context-dependence, I selected studies that reported mating behaviour in more than one environmental context. I focused on three mating behaviours: a) the amount of sexual signalling, the strength of response to mates or sexual stimuli (responsiveness), and the strength of mate choice (choosiness). I examined these behaviours in relation to seven social, biological or physical environmental factors: population density, adult sex ratio, operational sex ratio (OSR), predation risk, travel cost, time cost, and variation in mate quality. All of these factors potentially influence the costs and benefits of sexual signalling, mate searching or mate choice. They do this by altering several key components of the mating system: the number of potential mating opportunities, the cost of signalling, the cost of sampling, and the benefit of choice (**Table 1**). Importantly, as much as possible I avoid environmental factors which are likely to influence individual condition, because this is predicted to influence mating behaviour independently of the external environment (Cotton et al. 2006). This rules out other physical factors such as temperature or resource availability, which have the potential to influence both individual condition and some of the mating system components mentioned above.

**Table 1.** Outline of the key ways in which the seven environmental factors included in the meta-analysis have the potential to influence the expression of mating behaviour.

| Environmental factor | Environment influences: |  |  |  |
| --- | --- | --- | --- | --- |
|  | Mating opportunities | Cost of searching | Cost of signalling | Benefits of choice |
| Population density | ✓ | ✓ | ✓ |  |
| Adult sex ratio | ✓ | ✓ | ✓ |  |
| Operational sex ratio | ✓ | ✓ | ✓ |  |
| Predation risk | ✓ | ✓ | ✓ |  |
| Travel cost | ✓ | ✓ |  |  |
| Time cost | ✓ | ✓ |  |  |
| Variation in mate quality |  |  |  | ✓ |

Using this dataset, I performed multiple phylogenetically-controlled meta-analyses quantifying the difference in animal mating behaviour across environmental contexts.

Importantly, because I was interested in examining the overall effect of the environment on the expression of mating behaviour, I combined all seven environmental factors into a single analysis. However, I performed separate analyses for each of the three behaviours, as they are predicted to be influenced by the environment in different ways (see Predictions). I used this analysis to ask three questions. First, does sexual signalling, responsiveness and choosiness significantly differ across the animal kingdom in relation to the environment? Do animals respond in a consistent way, as would be expected from sexual selection theory?

Second, does the magnitude of this difference depend on which aspect of the environment is varied? Finally, are there any other aspects of the species tested, or experimental design used, that influence the direction or magnitude of this difference?

## Methods

### LITERATURE SEARCHES

I searched for relevant papers in two ways. First, I obtained all papers cited by a recent review of behavioural plasticity in mating behaviour (Ah-King & Gowaty 2016). Second, I performed literature searches using the online databases Web of Science & Scopus on the 29^th^ October 2018. The literature screening process is summarised in **Figure 1**. After removing duplicate results, I screened all titles to remove obviously irrelevant studies (e.g. studies on humans, other subject areas, review articles). I next imported all relevant abstracts into the screening software Rayyan (Ouzzani *et al*. 2016), and excluded those that did not appear relevant. This resulted in 701 relevant studies. I then read the full text of these 701 studies to determine if they fit the inclusion criteria listed below.

**Figure 1.**
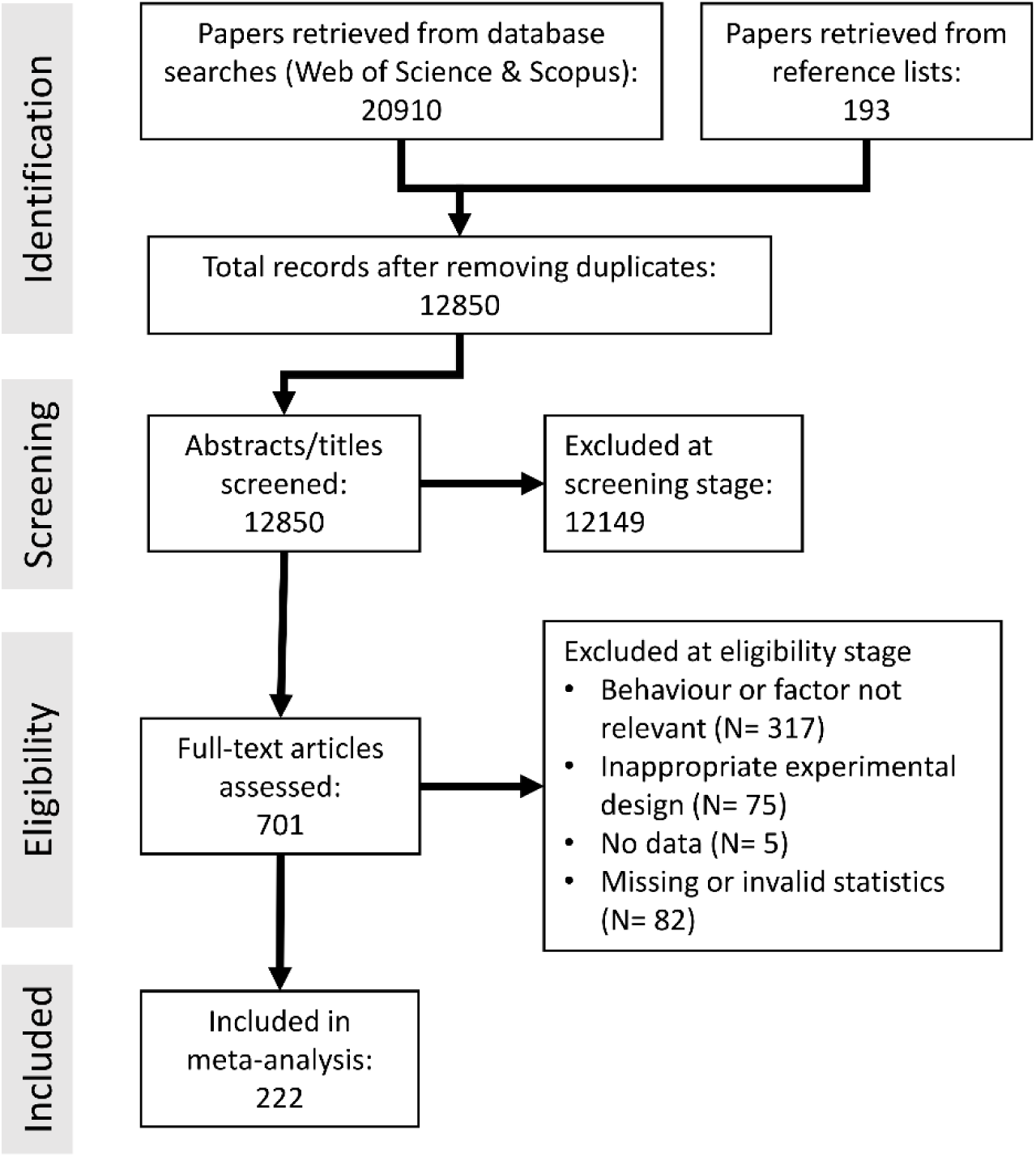
PRISMA diagram showing the literature search and selection process used to create the dataset.

### CRITERIA FOR INCLUSION

I had several main criteria for including a study in the next stage of the analysis. Studies were included that: a) measured one of the three mating behaviours listed above, b) recorded this behaviour in more than one context (in relation to one of the seven environmental factors listed above), and c) provided sufficient statistical information for an effect size to be calculated (see Effect size extraction and coding). I considered studies examining all animal species, with the exception of humans. I included studies testing the same subjects in multiple contexts, or different subjects in different contexts. I included data on both males and females, and from both experimental studies, in which environmental factors were directly manipulated, and observational studies examining behaviour in response to natural environmental variation. Importantly, I only included studies that could convincingly show that a single environmental factor was varied at a time. I did not include cases for which mating behaviour was inferred from mating outcomes (such as studies reporting metrics of sexual selection or mating frequency using paternity tests), or in which behaviour could not be attributed to a single individual (studies for which rivals or mates have some control over mating outcomes). I included studies in which subjects experienced a variable environment before or during the behavioural test. In the former case, the environment typically varied in the short term (hours or days before the trial), and so any responses seen can be considered to represent short-term behavioural plasticity. In a minority of cases, the environment was varied over a longer time period. For example, subjects may have been reared under different experimental conditions in the lab for several weeks. Some studies also compared the behaviour of wild-caught subjects from populations that differed naturally in environmental conditions.

### MATING BEHAVIOURS AND ENVIRONMENTAL FACTORS

Here, I briefly outline the inclusion criteria and predictions associated with the three behaviours and seven environmental factors included in the analysis. For a more detailed description of inclusion criteria and category definitions please see the supplementary methods.

I focused on three mating behaviours: sexual signalling, response to sexual signals (responsiveness), and the strength of mate choice (choosiness). In the sexual signalling category I included any behaviours (i.e. excluding morphological signals) that the authors suggest function to advertise to or attract mates. I included both long-range attraction signals (such as song or pheromone production, when mates are not immediately present), and close-range courtship behaviours that are expressed exclusively during mating interactions. Importantly, signalling behaviour was instead classed as choosiness if it was shown to be preferentially directed towards specific mates or phenotypes. I excluded non-behavioural signals (e.g morphology or colouration), or cases where it was unclear whether a signal had an exclusive sexual function (for example, male contest signals that are also used by females to assess males). Responsiveness can be defined broadly as the motivation to mate, or more strictly as the average response of an individual to potential mates or sexual signals (Brooks & Endler, 2001; Edward, 2015). A highly responsive individual is one that shows the strongest behavioural response across all presented mates or sexual stimuli. Responsiveness is a measure of the overall motivation to interact with potential mates or sexual stimuli, ignoring differences between options. In this category I included any mating behaviour (with the exception of sexual signalling, see above) summed or averaged across all options presented during a test. When such behaviours could be shown to be directed towards any specific mate, or type of mate, it was instead classed as choosiness (see supplementary material for more details). Choosiness is a measure of the strength of mate choice, which I define following Reinhold & Schielzeth (2015) as “the change in mating propensity in response to alternative stimuli”. In other words, the larger the difference in response to different stimuli, the choosier an individual is. In this category I included any mating behaviour for which the *difference* in response was compared between choice options. The greater the difference in response to sexual stimuli, the choosier the focal individual. The choosiness category includes any behavioural measure that can be interpreted as reflecting the strength of a mating preference. Preferences may be linked explicitly to a trait (either a specific stimulus or a mate phenotype), but this was not required for inclusion.

I focused on seven environmental factors: population density, adult sex ratio, operational sex ratio (OSR), predation risk, travel cost, time cost, and variation in mate quality (**Table 1**). The three social factors: density of conspecifics, adult sex ratio and OSR of the population, all provide information on the current number of available mating opportunities (Kvarnemo & Ahnesjo 1996; Kokko & Rankin 2006). The OSR is the ratio of reproductively active males to females in a population (Kvarnemo & Ahnesjo 1996), and so is the most salient piece of demographic information regarding current mating opportunities. Both the population density and adult sex ratio are imperfect measures of reproductive competition, but are much easier to assess. These factors also influence the amount of intrasexual competition, which could influence the payoffs associated with different mating tactics (Gross 1996; Weir *et al*. 2011). Finally, population density may also indirectly influence individual predation risk (Krause & Ruxton 2002). The population density category consists of studies comparing mating behaviour at different population densities, while controlling for the sex ratio perceived by subjects. In most cases, the sex ratio was equal (1:1). Importantly, I did not include cases in which population density could influence the amount of resources available to subjects, as this could potentially influence individual condition (Cotton *et al*. 2006).

I included one factor related to the biological environment: predation risk. For signallers, the risk of predation could influence the cost of conspicuous signalling and of searching for and sampling mates (Magnhagen 1991; Jennions & Petrie 1997; Zuk & Kolluru 1998). The level of predation may also influence the expected number of future mating opportunities via its effect on the density of conspecifics and the average expected lifespan (Hubbell & Johnson 1987; Ah-King & Gowaty 2016). I considered studies which tested both direct and indirect risk factors. Parasitoids can be considered ecologically similar to predators, because they lead to the death of the host, and so I also included studies examining the risk of parasitism by parasitoids in this category (but not studies examining the effect of other forms of parasitism).

I also included two factors relating to the physical environment: travel cost and time cost. The travel cost is the energetic cost (but not mortality cost) associated with movement, which should influence the cost of searching for and sampling mates (Real 1990: Jennions & Petrie 1997). The time cost is the amount of time remaining in the current breeding bout or mating season (Sullivan 1994), which influences the expected number of future mating opportunities, at least for the current season (Jennions & Petrie 1997). There is also the potential for other aspects of the environment to vary according to the season (such as population density or sex ratio), and so I only included studies in this category if the time of year was not explicitly linked to any other relevant environmental factors. I only included studies examining short-term time costs, rather than long-term changes associated with animal age, as this time cost may be confounded with other state-dependent effects when comparing individuals of different ages (Cotton *et al*. 2006).

Lastly, the variation in mate quality is the variation in mate phenotype experienced by the chooser, which is assumed to reflect variation in the direct or indirect benefits that will be received from mating with those individuals. Theory suggests that the benefits of being choosy are higher when mates vary greatly in quality (Parker 1983; Real 1990). For the variation in mate quality category, I excluded studies that did not control for the average mate quality experienced by subjects. This category only applies to choosiness and responsiveness.

There are other environmental factors that may influence mating behaviour in systematic ways that I did not consider here, because they do not influence the costs and benefits of expressing mating behaviour. For example, differences in noise or light levels instead reduce the ability of animals to *detect* or *discriminate between* signals (e.g. Seehausen *et al*. 1997; Swaddle & Page 2007; Candolin 2019), with the exception of studies that tested choosiness once a stimuli had been located. Additionally, other environmental stressors such as temperature may influence the costs and benefits of expressing mating behaviour (Candolin 2019), but these may also influence individual state. For example, in high-stress environments individuals may have less energy reserves to spend on costly mating behaviours (Coomes *et al*. 2019). I chose to exclude these types of stressors from the analysis, as there is no way of determining whether any behavioural change is driven by a context or state-dependent effect. For all three behaviours I excluded studies examining social experience effects that do not clearly influence the costs and benefits of choice, such as adult mate choice influenced by the phenotypes of parents or opposite-sex individuals encountered during development.

### PREDICTIONS

I predicted that choosiness should be highest, and so individuals should mate least randomly, when mating opportunities are common and the cost of sampling mates is low (low costs of choice), and when there is large variation in mate quality (high benefits of choice). Because of how I coded effect sizes (see Effect size extraction and coding), these predictions will result in a positive average effect for choosiness for all environmental factors (**Figure 2**).

**Figure 2.**
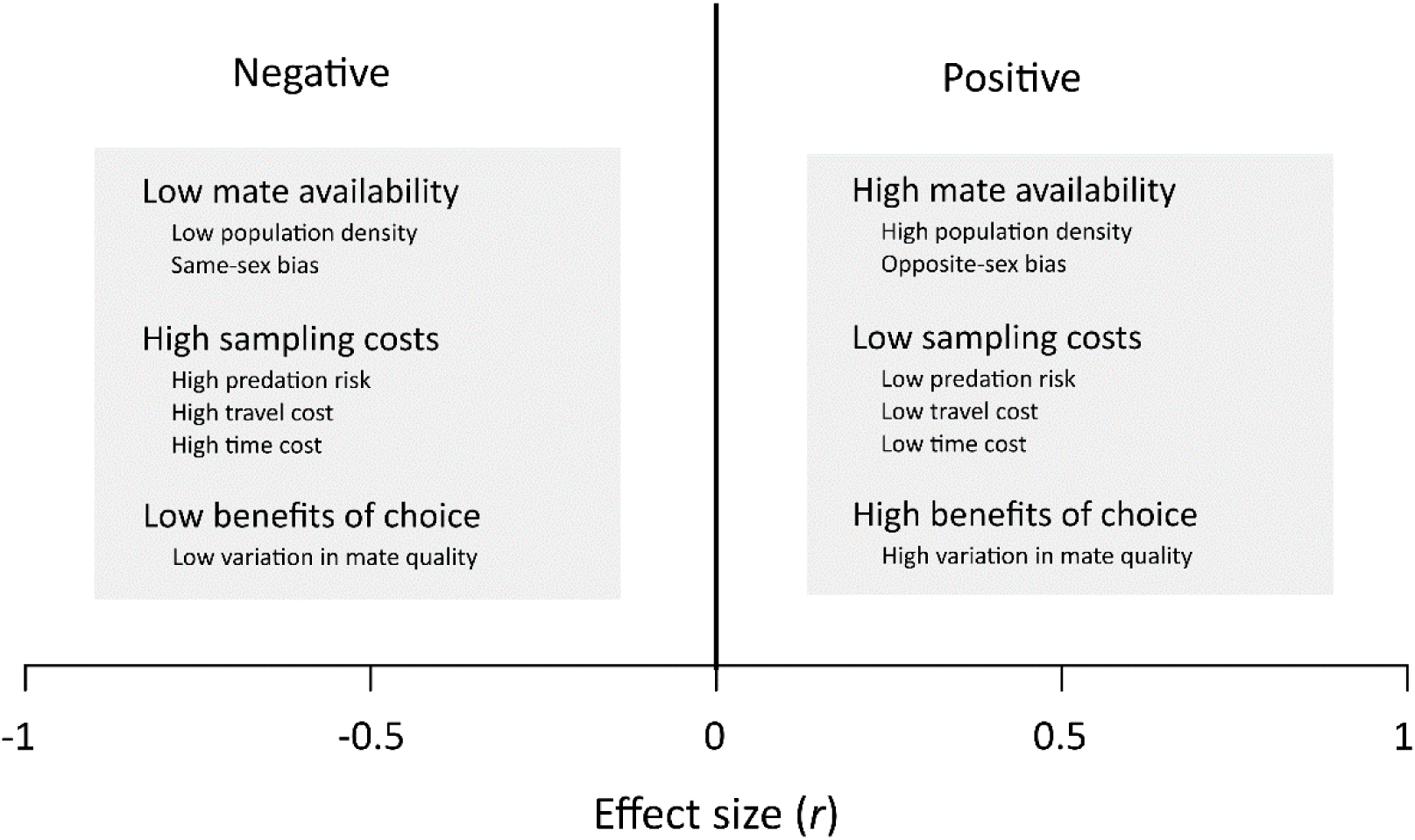
Diagram illustrating how differences in mating behaviour were assigned a positive or negative direction (in terms of the correlation coefficient *r*) in relation to environmental conditions. Positive effect sizes were assigned when mating behaviour was stronger under conditions of high mate availability, low costs of sampling mates and high benefits of mate choice. Negative effect sizes were assigned when mating behaviour was stronger under conditions of low mate availability, high costs of sampling mates and low benefits of mate choice. In all cases, ‘high’ and ‘low’ are relative terms, because environmental conditions were not standardised across studies.

The predictions for sexual signalling and responsiveness are less clear, because several processes could select for contrasting behavioural responses. If mate availability is most important for determining signalling and responsiveness, then sexual signalling and responsiveness should be highest when mating opportunities are rare and the cost of mate sampling is high, because in these situations each mating opportunity is potentially more valuable. This type of response is analogous to the ‘terminal investment’ observed in old or poor-condition individuals (Duffield et al. 2017). Alternatively, if signalling and mate searching are moderately costly, then individuals could conserve energy by reducing investment into these behaviours when the chances of securing a mate are low. Further, because signalling and mate searching generally increase predation risk, the expression of these behaviours may be greatest at a low predation risk (low cost of choice), as with choosiness (Zuk & Kolluru 1998). Finally, plasticity in sexual signalling and responsiveness could depend on the behaviour of chooser. If the more discriminating sex becomes choosier when mate availability is high, then courters will need to invest more into signalling and searching in these contexts in order to ensure a mating. Therefore, depending on which processes are most important, the average effect size for sexual signalling and responsiveness could be negative (if mate availability is most important) or positive (if conserving available energy reserves or responding to choosers is most important) (**Figure 2**).

### EFFECT SIZE EXTRACTION AND CODING

I used the correlation coefficient *r* as the measure of effect size. In this analysis, the effect size represents the *difference* or *change* in a behaviour due to the environment. Larger values therefore represent a greater difference in behaviour across different contexts. An effect size of zero indicates no difference in behaviour across contexts, but is blind to the magnitude across contexts. For all analyses, I used Fisher’s *Z* transform of the correlation coefficient (*Zr*), as r is constrained within ± 1 and so does not adhere to Gaussian distribution assumption (Koricheva *et al*. 2013). The associated variance for *Zr* (var *Z*) was calculated as 1/(n - 3) (Borenstein *et al*. 2009), with n being the total number of animals used in the test.

I extracted all relevant effect sizes from each study. In many cases this resulted in multiple effect sizes per study, because studies often report results from multiple experiments, or compare several different behaviours measured in the same experiment. The potential non- independence arising from using multiple effect sizes per study is controlled for in the statistical analysis (see Statistical Analyses). In many cases I obtained measurements for more than one behavioural category from a single study (though note that I ran separate analyses for each category). When statistical information was available, I obtained effect sizes either directly (if correlations were reported in the text), using summary data presented in the study, or from the results of statistical tests, using a range of conversion equations (Lipsey & Wilson 2001; Koricheva *et al*. 2013). I used two approaches to obtain effect sizes when appropriate statistics were missing. First, where possible I performed my own analyses using reported summary statistics, or raw data, presented in the text, in tables and figures, or in available supplementary results or data. I used the online tool WebPlotDigitizer v4 (Rohatgi 2019) to extract raw data from scatter plots, and means and standard deviations from bar plots. Second, I contacted authors directly and asked for either summary statistics or raw data. I obtained data this way for 17 studies (Berglund 1994; Evans & Magurran 1999; Evans *et al*. 2002; Velez & Brockmann 2006; Wong & Svensson 2009; Young *et al*. 2009; Ziege *et al*. 2009; Lafaille *et al*. 2010; Makowicz *et al*. 2010; Willis *et al*. 2012; Pilakouta & Alonzo 2013; Franklin *et al*. 2014; Wilgers *et al*. 2014; Breedveld & Fitze 2015; Pompilio *et al*. 2016; Filice & Long 2017; Pilakouta *et al*. 2017).

The original direction of the extracted effect sizes is not meaningful, as it depends on the type of data used (for example: association time is positively related to preference, whereas approach latency is negatively related to preference), or which treatment is classed as the control. I therefore manually assigned a direction to all effect sizes, in relation to the environmental context under which behaviours were more strongly expressed. I assigned directions based on the hypothesised costs of mate searching and mate choice (but not sexual signalling). I assign a positive direction to conditions in which the cost of expressing mate searching and mate choice is expected to be low. This is associated with high mate availability and low energetic or mortality costs of mate sampling (**Figure 2**). Conversely, I assign a negative direction to conditions in which the cost of mate searching and mate choice is expected to be high, so that each mating encounter is more valuable. Therefore, the effect size was assigned a positive direction when sexual signalling, responsiveness or choosiness was highest when: the population density is high, the adult sex ratio or OSR is biased towards the other sex, the predation risk is low, the travel and time costs are low, and there is large variation in mate quality (**Figure 2**). Conversely, the effect size was assigned a negative direction when sexual signalling, responsiveness or choosiness was highest when: the population density is low, the adult sex ratio or OSR is biased towards the same sex, the predation risk is high, the travel and time costs are high, and there is small variation in mate quality (**Figure 2**). I note also that the terms ‘high’ and ‘low’ in this case are relative, because the actual environmental conditions are not standardised across studies. So the phrase ‘high predation risk’ is shorthand for ‘the context in which predation risk in highest’.

In several cases, studies presented tests statistics that were non-significant, but provided no descriptive or statistical information that allowed me to determine the direction of an effect (for example, chi-squared statistics do not encode which cell has the larger value). These effect sizes would traditionally not be included in a meta-analysis in which effect size direction is important. However, this systematically biases the dataset against non-significant results (Harts *et al*. 2016), as such information is almost always available for significant results, for which the direction of an effect is seen to be more important. As a form of sensitivity analysis I assumed that these effect sizes were equally likely to be weakly positive or negative, and assigned them a value of zero. I then ran the analyses with and without including these directionless data points. This process resulted in six separate datasets: a zeros included dataset and a zeros excluded dataset for each behaviour category.

### PHYLOGENETIC TREES

In order to control for the potential non-independence of effect sizes due to shared evolutionary history (Hadfield & Nakagawa 2010; Koricheva *et al*. 2013) I created a phylogeny of the species included in each of the six datasets. Given the broad range of species included in each sample, no single published phylogeny was available that included all species. I therefore constructed a phylogenetic supertree for each of the six datasets using the Open Tree of Life (OTL) database (Hinchliff *et al*. 2015) and the rotl R package (Michonneau *et al*. 2019). Given the absence of accurate branch length data for these trees, all branch lengths were therefore first set to one and then made ultrametric using Grafen’s method (Grafen 1989), using the R package ape v5.3 (Analyses of Phylogenetics and Evolution: Paradis *et al*. 2014). In cases where the OTL database resulted in a polytomy, I manually searched for published phylogenies that could resolve them (see supplementary methods for details). The final ultrametric trees for the three full datasets (zeroes included) can be seen in the supplementary material (**Figures S1-S3**).

### MODERATORS

I tested for the effect of 10 categorical moderator variables (eight for each behaviour) on the size or direction of context-dependent plasticity.

#### Environmental factor

The predicted directions for each factor are related to the way effect sizes were coded, and are described above (see Predictions). Additionally, for all behaviours I predicted that predation risk should induce the greatest degree of plasticity, due to the strong fitness consequences of failing to respond to this risk.

#### Focal sex

I compared the strength of plasticity between males and females. Any effect sizes for which the sex of the focal subjects was not specified were classed as ‘both’. I predicted that, for all behaviours, males should show reduced plasticity compared to females. This is because males typically have more to gain from mating often, and so are more likely to invest highly in mating across multiple contexts. Additionally, the fact that males typically show weaker mate choice than females means they are able to reduce choosiness to a lesser extent compared to females, potentially resulting in lower apparent plasticity in choosiness for males.

#### Taxonomic class

I assigned species to taxonomic groups at the class level. I predicted that, for all behaviours, plasticity should be smallest for short-lived species such as insects. This is because immediate reproductive success is more important for short-lived species, which cannot afford to wait for environmental conditions to change to the same extent that longer-lived species can.

#### Environmental factor timing

I compared studies in which subjects were exposed to environmental differences during a behavioural trial to those in which subjects were exposed to environmental differences before a trial (and in some cases the environment was varied both before and during a trial). Predictions for how this moderator would influence the strength of plasticity were less clear. Given that behavioural trials are typically short, before-test environmental differences may result in more plasticity, as subjects experience this environment for longer. However, some factors, especially predation risk, are more relevant at the exact moment the behaviour is being expressed, especially in highly heterogeneous environments.

#### Environmental factor variation

I compared studies in which environmental factors were manipulated to those which examined natural environmental variation. I predicted that behavioural plasticity should be stronger in experimental tests, as environmental differences are likely to be more consistent in these studies.

#### Animal origin

I classed focal subjects as either lab-reared, wild-caught (and observed in the laboratory) or wild (and observed in the field). I predicted that behavioural plasticity should be greatest in lab-reared individuals, because the experience of potentially-confounding effects may be reduced in a laboratory environment.

#### Signalling modality (sexual signalling only)

Sexual signalling was classed as either visual, acoustic, chemical, tactile, or mixed (if the signal was multi-modal and specific behavioural components were not recorded). I predicted that plasticity would be greatest for visual and acoustic signals, as these have the potential to be costlier than chemical or tactile signals, both in terms of energetic investment and the risk of being intercepted by predators.

#### Signalling type (sexual signalling only)

I recorded whether signals were produced at close-range (when potential mates or other sexual stimuli were present) or long-range (when no mates or sexual stimuli were present). I predicted that plasticity should be greatest for long-range signalling, as this is likely to be costlier than short-range signalling (both in terms of energetic investment and conspicuousness to predators), and so should only be expressed in favourable environments.

#### Preference measure (responsiveness and choosiness)

I recorded whether responsiveness or choosiness were determined using actual mating events, or a ‘proxy’ behavioural measure of preference that is assumed to correlate with mating (Dougherty 2020). Proxy measures of preference include association time, approach latency, and male-female interactions prior to mating. I predicted that studies recording mating would show reduced plasticity compared to proxy behaviours. This is because mating results from an interaction between two individuals. Therefore, any misalignment in experience or motivation between the two parties will reduce the ability of either individual to show strategic context-dependent behaviour.

#### Stimuli type (responsiveness and choosiness)

I recorded whether subjects were presented with conspecific signals only (‘mate-quality tests’), or could choose between conspecific and heterospecific signals (‘species recognition tests’). I predicted that subjects would show greater plasticity in mate-quality tests, because subjects in species recognition tests should prefer conspecifics in almost all environments (but see Willis 2013).

### STATISTICAL ANALYSES

All statistical analyses were performed using R v3.6 (R development Core Team 2019). Meta-analyses were performed primarily using the meta-analysis package Metafor v2.1 (Viechtbauer 2010). In order to determine the overall mean effect size for each dataset, I ran a multilevel random-effects model using the rma.mv function, with study, species, and phylogeny as random factors (Nakagawa & Santos 2012). Phylogeny was incorporated into the model using a variance-covariance matrix, assuming that traits evolve via Brownian motion. The Fisher’s *Z* transformation was used as the effect size in all models, and model results were then converted back to *r* for presentation. The mean effect size was considered to be significantly different from zero if the 95% confidence intervals did not overlap zero. I ran these overall models separately for each of the three behaviours. For each behaviour, I ran models with and without the inclusion of effect sizes of no direction (all effect sizes set at zero).

I used I^2^ as a measure of heterogeneity of effect sizes (Higgins *et al*. 2003). I^2^ values of 25, 50 and 75% are considered low, moderate and high, respectively (Higgins *et al*. 2003). I calculated I^2^ across all effect sizes, and also partitioned at different levels of the model using the method of Nakagawa & Santos (2012). This allowed me to quantify the amount of variation in effect size that could be attributed to differences in study, species, and phylogenetic history.

I detected significant heterogeneity in effect sizes for all three behaviours, and so I therefore investigated potential moderators of effect size using the full dataset for each behaviour (i.e. all zeroes included). To test for the effect of moderators I ran meta-regression models, which were identical to the above models except for the inclusion of a single categorical or continuous fixed factor. I ran a separate model for each fixed effect. I considered a moderator to significantly influence the mean effect size by examining the *Q*_*M*_ statistic, which performs an omnibus test of all model coefficients. For each behaviour I tested the effect of nine moderators: eight categorical and one continuous (study year). I tested the effect of different moderator variables depending on the behaviour examined. In order to estimate the average effect size for each level of a categorical factor I ran the same meta-regressions as above, but excluding the model intercept (again run separately for each fixed factor). For sexual signalling and responsiveness the sample sizes for some environmental factors were small. Therefore, in order to check the sensitivity of the meta-regressions testing the effect of environmental factor, I ran each of these tests twice for each behaviour: first including all factors, and second after removing any categories with 6 or less effect sizes.

Finally, I searched for signs of two types of publication bias in the three (zeroes included) datasets. I first searched for signs of time-lag bias, which arises when earlier published studies have larger effect sizes than later published studies, which may indicate bias against publishing studies of small effect in young research fields (Koricheva *et al*. 2013). To test for any change in effect size over time, I ran a meta-regression with study year as a fixed effect (mentioned above). Second, I searched for signs of publication bias against studies with small sample sizes or non-significant results (Koricheva *et al*. 2013), by looking for funnel plot asymmetry using a trim-and-fill test (Duval & Tweedie 2000) and Egger’s regression (regression of *Zr* against inverse standard error; Egger *et al*. 1997).

## Results

### SEXUAL SIGNALLING

I obtained 260 effect sizes examining context-dependent sexual signalling, from 114 studies and 68 species. I obtained data from seven taxonomic groups, though the majority of studies considered insects and fish (**Figure 3a**). The majority of effect sizes considered male signalling (males: k= 230; females: k= 24; no sex specified: k= 6).

**Figure 3.**
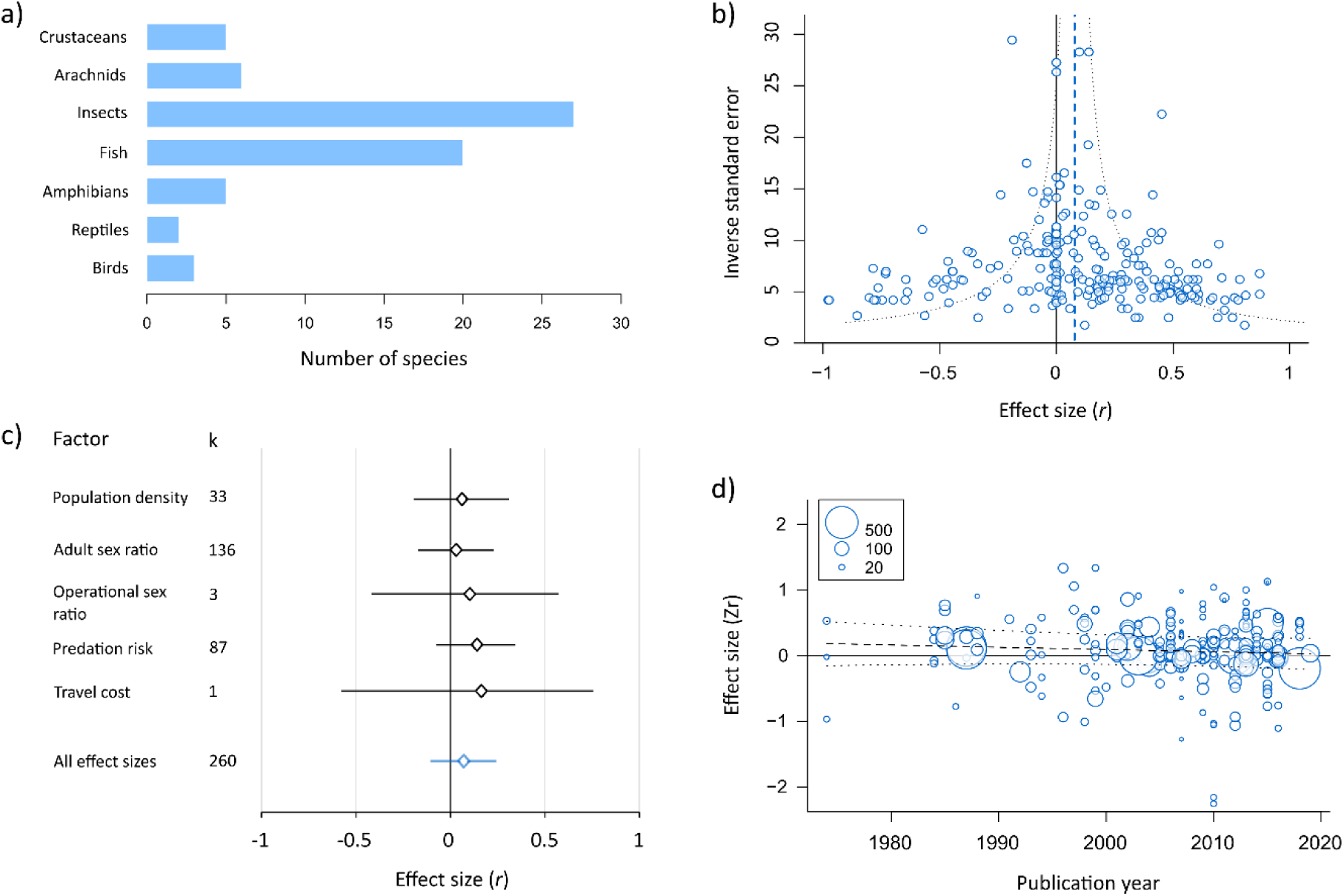
Summary results for context-dependent sexual signalling. a) Histogram showing the number of species included in relation to taxonomic grouping. b) Funnel plot showing the relationship between effect size (r) and sample size (inverse standard error). The dotted line shows the mean effect size for the full model. c) Forest plot showing the average effect size for each environmental factor separately. In all cases diamonds represent the mean effect size estimate, and the bars represent the 95% confidence interval. The mean effect size obtained from the full model, across all effect sizes, is shown in blue for comparison. k is the number of effect sizes in each category. d) Bubble plot showing the relationship between effect size (Zr) and publication year. The points are scaled by the sample size of each estimate. The broken line shows the predicted regression line from a meta-regression, and the dotted lines are the 95% confidence intervals.

Overall, sexual signalling behaviour did not consistently differ across contexts, either using the full dataset (k= 260, mean= 0.07, 95% CI= -0.11-0.24 **Figure 3b**) or the reduced dataset (k= 209, mean= 0.095, 95% CI= -0.12-0.18). The full dataset shows very high heterogeneity across effect sizes (Total I^2^= 93.4%), with 36.4% being attributable to between-study differences, <1% to between-species differences, and 11.24% to phylogenetic history.

Given the high heterogeneity across effect sizes, I tested whether this variation could be explained by differences in species traits or study methodology. The strength or direction of the signalling response did not differ for the five environmental factors tested (**Table 2**; **Figure 3c**): for all environmental factors signalling was greatest when the cost of choice was low (positive effect size), however the mean effect size did not differ from zero for any environmental factor individually. This result remained after removing the two environmental factors with 6 or less effect sizes (OSR and travel cost, Q_M 2_= 2.33, P= 0.31, k= 256). The average signalling response did not differ according to any of the other moderators tested, including taxonomic class or focal sex (**Table 2; Table S1**).

**Table 2.** Meta-regression results for all three behaviours. Significance was determined using a *Q*_*M*_ test for both categorical and continuous fixed effects. Each factor was tested using a separate mixed-effects model, with a single fixed factor and four random factors (Study ID, species, phylogeny and observation ID). Significant factors are highlighted in grey.

| Fixed effect | Signalling |  | Responsiveness |  | Choosiness |  |
| --- | --- | --- | --- | --- | --- | --- |
| | $Q_M$ | $P$ | $Q_M$ | $P$ | $Q_M$ | $P$ |
| Environmental factor | 2.44 | 0.66 | 9.50 | 0.09 | 8.89 | 0.18 |
| Focal sex | 1.08 | 0.58 | 0.85 | 0.65 | 5.40 | 0.07 |
| Taxonomic class | 2.19 | 0.9 | 2.44 | 0.93 | 3.33 | 0.85 |
| Factor timing (Before vs during test) | 2.78 | 0.25 | 3.48 | 0.18 | 0.39 | 0.82 |
| Factor variation (Manipulated vs natural) | 1.09 | 0.3 | 0.01 | 0.93 | 0.01 | 0.93 |
| Animal origin (Wild vs lab-reared) | 0.42 | 0.81 | 3.64 | 0.16 | 1.81 | 0.61 |
| Signalling modality | 2.74 | 0.6 | - | - | - | - |
| Signalling type (Short vs long range) | 0.04 | 0.84 | - | - | - | - |
| Preference measure (Matings vs proxy) | - | - | 0.20 | 0.66 | 0.14 | 0.70 |
| Stimuli type (Mate-quality vs species recognition) | - | - | 0.07 | 0.79 | 1.37 | 0.24 |
| Study year | 0.78 | 0.38 | 0.001 | 0.98 | 8.78 | 0.003 |

### RESPONSIVENESS

I obtained 176 effect sizes examining context-dependent differences in responsiveness, from 86 studies and 53 species. I obtained data from eight taxonomic groups, though the majority of studies considered insects and fish (**Figure 4a**). I obtained an approximately equal number of effect sizes from both sexes (males: k= 78; females: k= 80; no sex specified: k= 18).

**Figure 4.**
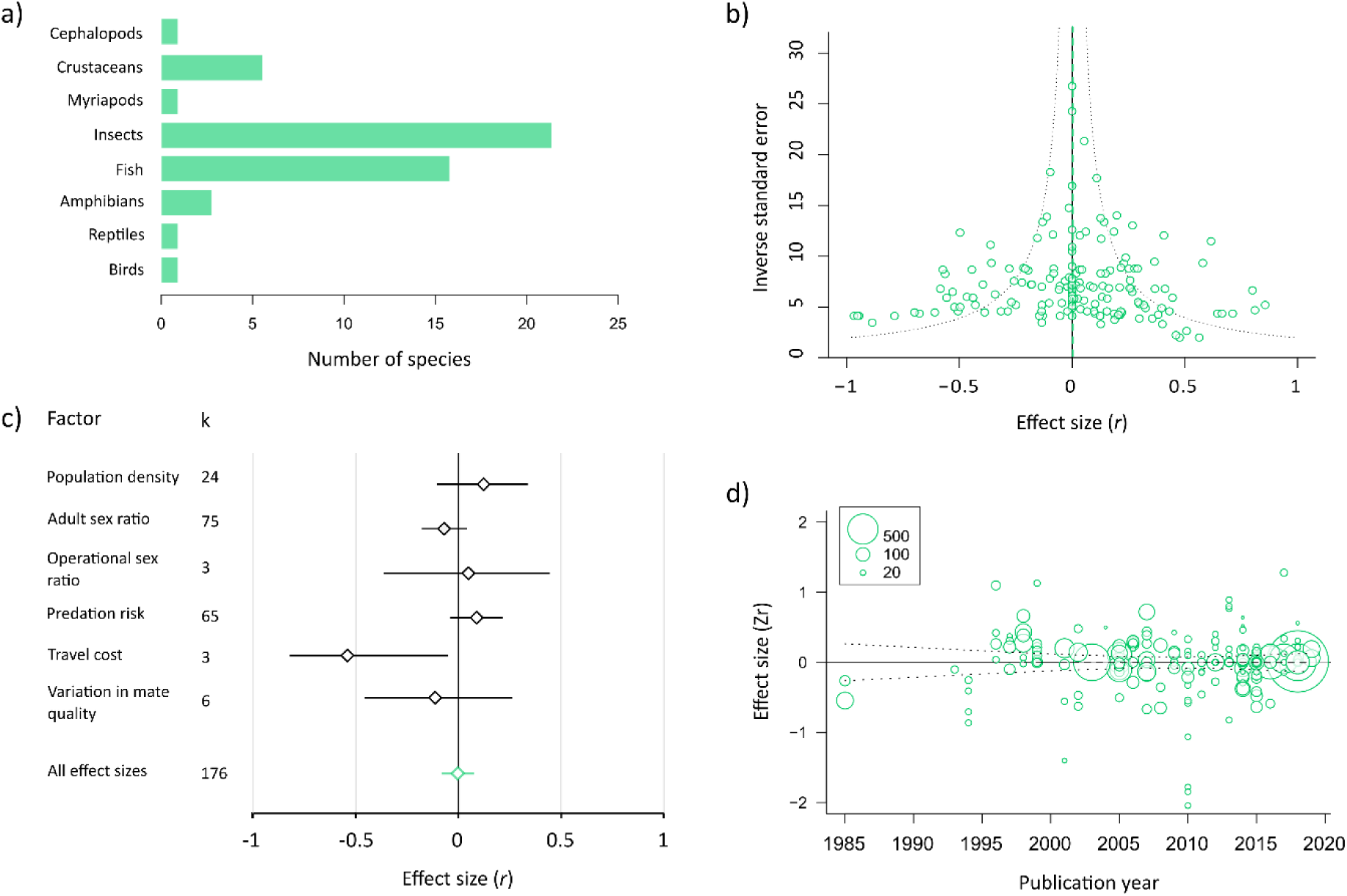
Summary results for context-dependent responsiveness. a) Histogram showing the number of species included in relation to taxonomic grouping. b) Funnel plot showing the relationship between effect size (r) and sample size (inverse standard error). The dotted line shows the mean effect size for the full model. c) Forest plot showing the average effect size for each environmental factor separately. In all cases diamonds represent the mean effect size estimate, and the bars represent the 95% confidence interval. The mean effect size obtained from the full model, across all effect sizes, is shown in green for comparison. k is the number of effect sizes in each category. d) Bubble plot showing the relationship between effect size (Zr) and publication year. The points are scaled by the sample size of each estimate. The broken line shows the predicted regression line from a meta-regression, and the dotted lines are the 95% confidence intervals.

Overall responsiveness did not consistently differ across contexts, either using the full dataset (k= 176, mean= -0.003, 95% CI= -0.082-0.08; **Figure 4b**) or the reduced dataset (k= 146, mean= -0.001, 95% CI= -0.1-0.1). The full dataset shows very high heterogeneity across effect sizes (Total I^2^= 91.6%), with 67.5% being attributable to between-study differences, and <1% to between-species differences or phylogenetic history.

The difference in responsiveness was not significantly influenced by environmental factor (**Table 2**). This result remained after removing the three environmental factors with 6 or less effect sizes (OSR, travel cost and variation in mate quality, Q_M 2_= 4.51, P= 0.11, k= 164). In the full model there was a tendency for a positive effect size for predation risk, population density and OSR and a negative effect size for adult sex ratio, travel cost and variation in quality (**Figure 4c**). The average difference in responsiveness was not significantly influenced by any of the other moderators tested (**Table 2; Table S2**).

### CHOOSINESS

I obtained 261 effect sizes examining context-dependent differences in choosiness, from 105 studies and 61 species. I obtained data from eight taxonomic groups, though the majority of studies considered insects and fish (**Figure 5a**). Female choice is more common than male choice in the dataset (female choice: k= 159; male choice: k= 96; no sex specified: k= 6).

**Figure 5.**
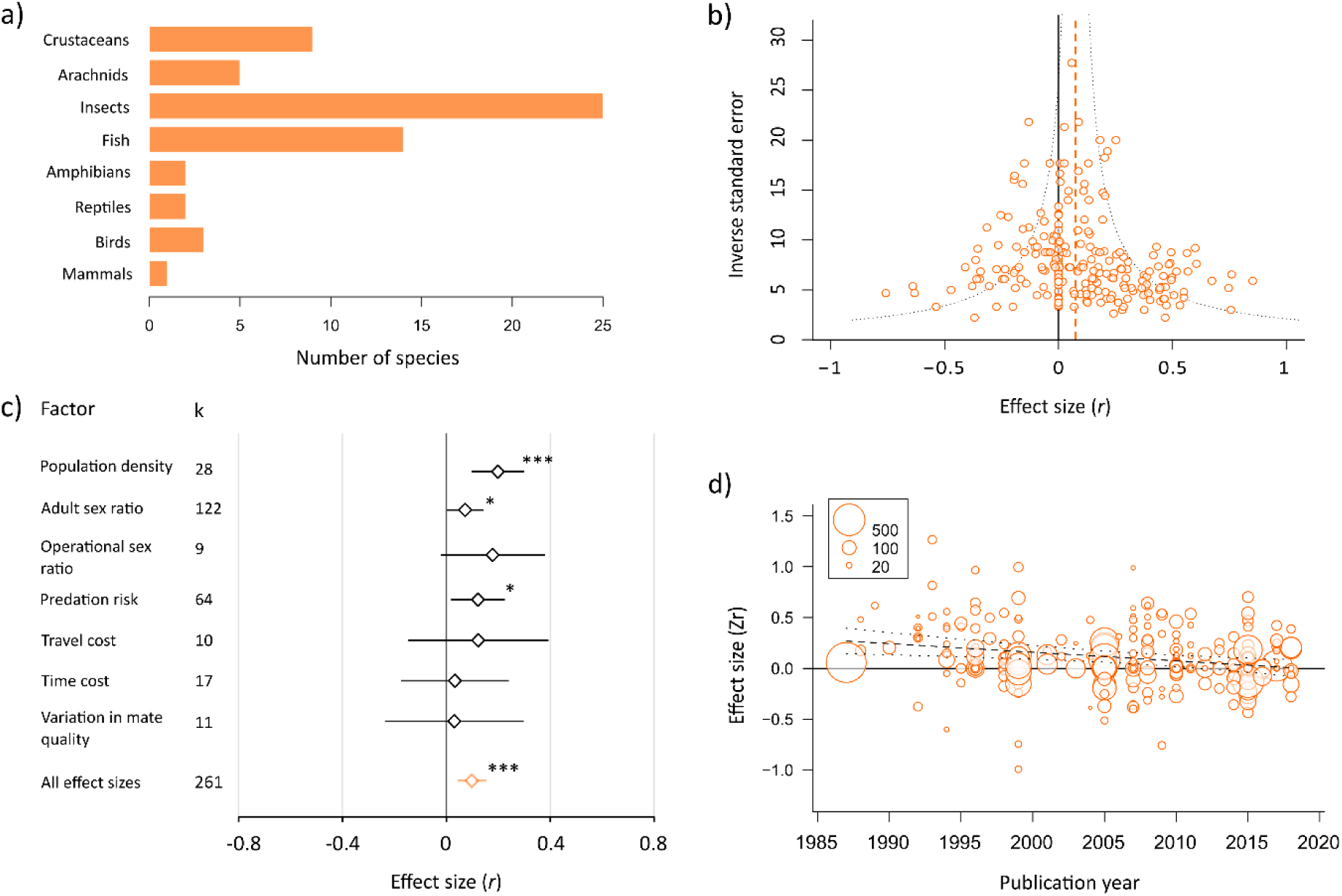
Summary results for context-dependent choosiness. a) Histogram showing the number of species included in relation to taxonomic grouping. b) Funnel plot showing the relationship between effect size (r) and sample size (inverse standard error). The dotted line shows the mean effect size for the full model. c) Forest plot showing the average effect size for each environmental factor separately. In all cases diamonds represent the mean effect size estimate, and the bars represent the 95% confidence interval. The mean effect size obtained from the full model, across all effect sizes, is shown in orange for comparison. k is the number of effect sizes in each category. Estimates that differ significantly from zero are marked with asterisks (*, P< 0.05; **, P< 0.01; ***, P < 0.001). Bubble plot showing the relationship between effect size (Zr) and publication year. The points are scaled by the sample size of each estimate. The broken line shows the predicted regression line from a meta-regression, and the dotted lines are the 95% confidence intervals.

Overall, choosiness was significantly higher when the costs of mate choice were low (Full dataset, k= 261, mean= 0.098, 95% CI= 0.043-0.16; **Figure 5b**). This result was the same after removing 65 effect sizes set at zero (reduced dataset, k= 196, mean= 0.12, 95% CI= 0.05-0.19). However, the overall effect size is small (Cohen 1992). The full dataset shows very high heterogeneity across effect sizes (Total I^2^= 81.2%), with 40.9% being attributable to between-study differences, 17.9% to between-species differences, and <1% to phylogenetic history.

The difference in choosiness across contexts was not significantly affected by environmental factor (**Table 2**); the average estimate was positive for all factors, but significantly differed from zero for predation risk, population density, and adult sex ratio (**Figure 5c**). The average estimates for operational sex ratio, travel cost, time cost and variation in mate quality did not differ significantly from zero, however all four categories consisted of a small number of effect sizes (k <20), so this lack of an effect should be interpreted with caution. The average choosiness response was not significantly influenced by any of the other moderators tested (**Table 2; Table S3**).

### PUBLICATION BIAS

Evidence for publication bias was mixed. The difference in signalling and responsiveness was not significantly related to study year (**Table 2; Figures 3d & 4d**). However, the average choosiness response decreased significantly over time (**Table 2**; **Figure 5d**). A trim-and-fill test did not detect any ‘missing’ effect sizes for choosiness. However, significant asymmetry was detected for responsiveness and sexual signalling, with 28 and 24 ‘missing’ negative effect sizes respectively (**Figure S4**). Inclusion of these effect sizes did not change the interpretation of the sexual signalling data (the overall mean was still not significantly different from zero: mean= 0.03, 95% CI= -0.02-0.07), but resulted in a significantly negative effect size for responsiveness (mean= -0.07, 95% CI= -0.12 -0.02). A regression test did not detect any significant relationship between effect size and study variance for sexual signalling (*F* _1, 258_ = 0.41, P= 0.52; **Figure S5**) or responsiveness (*F* _1, 174_= 0.19, P= 0.67; **Figure S5**), but there was a significant effect for choosiness (*F* _1, 259_= 4.87, *P*= 0.028; **Figure S5**). This latter effect seems to be driven by a lack of negative effect sizes of low power, which is suggestive of publication bias.

## Discussion

Investment in mating behaviour is often costly, and the fitness payoffs of this investment can vary across contexts. Therefore, animals are expected to alter their mating behaviour depending on the current context, in order to minimise the amount of investment needed to secure matings, and maximise fitness outcomes. By synthesising the results of 222 studies and 697 effect sizes examining animal mating behaviour across multiple contexts, I found that choosiness (the strength of mate choice) differed significantly across environments.

Choosiness was significantly stronger in contexts where the cost of mate choice is low, such as when mating opportunities are frequent and the perceived risk of predation is low. However, the average effect of each factor alone was much weaker than expected, and there was some evidence for a decrease in effect size over time. Neither sexual signalling nor responsiveness differed across contexts in a consistent way, either across the whole dataset or when each environmental factor was considered individually. Taken together, these results suggest that the expression of mate choice is more context-dependent than either sexual signalling or responsiveness, but that overall the evidence for context-dependent mating behaviour across animals is currently surprisingly weak. The common assumption that animal mating behaviour shows context-dependent expression may need to be reassessed in light of these findings.

Why might mate choice be more consistently sensitive to the environment than sexual signalling or responsiveness? One explanation is that the environmental factors examined here are predicted to influence choosiness in the same way: that is, when conditions become unfavourable, choosiness should decrease. In contrast, there may be conflicting selection pressures acting on signalling and responsiveness which cause the direction of plasticity to differ across species or contexts. For example, theory suggests the point at which switching to invest maximally in reproduction depends on the severity of the current environmental conditions, and the potential number of future mating opportunities an individual will have (Duffield et al. 2017). However, the problem with this explanation is that both of these factors are poorly accounted for in my models, so my ability to support it is limited (and I come back to the issue of explaining heterogeneity below). An alternative explanation relates to the relative importance of the behaviours for reproductive fitness.

While choosing the right partner can often provide strong fitness benefits to choosers (Andersson 1994; Kokko *et al*. 2003), the consequence of failing to express a choice is mating with a random partner, which will only be costly when most potential partners are of low quality. However, reduced signalling or mate searching may often lead to a complete failure to mate, resulting in a fitness of zero. Therefore, in many contexts gaining any mate (which may require investment in mate searching and/or sexual signalling) should be more important than gaining a *high-quality* mate. One consequence of this could be a mating strategy whereby investment in sexual signalling and mate searching is high under most conditions, which will result in reduced plasticity in response to the environment.

All three datasets were characterised by very high heterogeneity in both the strength and direction of the effect size. Sexual signalling and responsiveness in particular showed an approximately equal number of positive and negative effect sizes, including many of large magnitude. Partitioning of the model variances suggested that little heterogeneity could be explained by species differences or phylogenetic relatedness, and for all three behavioural types the majority of this heterogeneity remains unexplained. I therefore tested whether a range of biological and methodological moderating factors could explain this variation. Importantly, environmental factor, sex or taxonomic group did not significantly explain the variation in any behaviour (while choosiness was context-dependent, this response did not differ according to which environmental factor was examined). In fact, for sexual signalling and responsiveness, the mean effect size for each environmental factor considered alone did not differ significantly from zero. Choosiness was highest when the costs of choice were lower for all of the seven factors tested, though the mean effect size was significantly different from zero only for population density, adult sex ratio, and predation risk. However, the lack of a significant effect for travel cost, time cost and variation in mate quality are likely driven by the small sample sizes for these groups, and so any conclusions relating to these factors should be interpreted with caution. Interestingly, choosiness was more sensitive to differences in population density than to differences in sex ratio, even though the latter is a more accurate measure of the number of available mating opportunities.

Individuals may be more likely to respond to changes in overall population density if it is easier to assess accurately. Alternatively, this effect could be driven by the fact that population density tends to vary more than sex ratio in an absolute sense. For example, across all studies included in the three datasets that measured or manipulated population density (N= 22), the median number of conspecifics was 4 (±6.8) at low density and 20.5 (±56.3) at high density. Assuming a 1:1 sex ratio, this corresponds to 2 and 10 ‘available’ mates in these studies. In comparison, for studies that measured or manipulated sex ratio across all three datasets (N= 98), the median number of mates per focal individual is 0.5 (±1.4) at low mate availability and 2 (±8.3) at high mate availability.

Importantly, the majority of heterogeneity in all three datasets remained unexplained after testing the effects of ten moderating factors. It is unclear whether such heterogeneity represents real, biological variation or stems from some other source. Some of this variation is likely due to sampling error, as for all three behaviours many of the largest effect sizes are associated with studies with small sample sizes, and thus low statistical power. Some of this variation could also be explained by methodological limitations. For example, the effect size used here is only able to detect linear effects. This means that significant quadratic effects, such as peak signalling at intermediate densities (Kokko & Rankin 2006), will not be captured here. Alternatively, the large variation observed may be the result of methodological differences between studies that have not been accounted for (Dougherty & Shuker 2015; Rosenthal 2017; Dougherty 2020). For example, studies typically assume animals can accurately assess the costs of expressing a behaviour in a given environment, but this may not always be the case. Therefore, differences in the extent to which studies successfully manipulate these perceived costs may lead to significant variation in context-dependent behavioural responses. Experimental studies may also often use subjects that are especially eager to mate, for example because they are virgin or have been isolated from members of the opposite sex, and such individuals are predicted to show lower levels of context-dependent behaviour than mated or experienced individuals (Ah-King & Gowaty 2016; Kelly 2018). Finally, the observed heterogeneity may stem from biological differences that are difficult assess for all of the species sampled, for example in relation to mating system, life-history or the energetic costs of signalling. Importantly, one key factor that is currently unaccounted for is the cost of expressing mating behaviour in a given environment: plasticity should be largest where behaviours are compared across environments that differ greatly in the costs and benefits of expression. This is important, because the included studies differ in terms of the range of environmental conditions subjects are tested in (environmental differences are not standardised across studies), and so will differ also in the range of any environment-induced costs. Unfortunately, we simply do not have accurate data on what these costs are, even for a small number of behaviours or contexts. This is likely to be the case for some time, given the difficulty in measuring fitness in ecologically relevant contexts. However, without this data we also cannot rule out the possibility that experiments simply do not present subjects with a sufficiently variable range of contexts.

In conclusion, this study suggests that the evidence that animal mating behaviour varies in a consistent way across different environments is currently quite limited. Across species, sexual signalling and responsiveness do not appear to consistently respond to any of the environmental differences tested. Choosiness did show consistent, significant differences in relation to predation risk, population density and adult sex ratio, but the effect sizes are generally weaker than expected. This is despite plenty of good empirical examples of context-dependent mating behaviour as predicted by sexual selection theory, and narrative reviews consisting almost entirely of affirmatory examples (e.g. Ah-King & Gowaty 2016; Kelly 2018). Importantly, the datasets for all three behaviours were characterised by very high heterogeneity in effect size which remains mostly unexplained, even after accounting for the phylogenetic relationships amongst species. It therefore remains unclear whether environmental variability is a less important driver of behavioural plasticity than predicted, or whether the lack of a strong effect is due to unaccounted biological or ecological variability across species, or excessive ‘noise’ arising from methodological differences across studies or low statistical power. The best way to try to tease apart these alternatives in the future will be to perform careful, well-designed studies, preferably using large sample sizes. This work is needed if we are to understand the expression of animal mating behaviour, and evolutionary forces driven by mate choice and intrasexual competition, in complex and rapidly-changing natural environments. Further, human-induced changes in the natural environment have the potential to influence most of the factors considered here (e.g. population density, predator density, travel cost, time cost). Therefore, understanding how mating behaviour and population fitness respond to these increasingly challenging natural conditions will help us to predict whether natural populations will be able to adapt and persist in the wild.

## Acknowledgements

This work was funded by a Leverhulme Trust Early-Career Fellowship (ECF-2018-427). I would like to thank Anders Berglund, Jane Brockmann, Jonathan Evans, David Filice, Patrick Fitze, Amanda Franklin, Michael Greenfield, Lotta Kvarnemo, Amber Makowicz, Gabriel Manrique, Natalie Pilakouta, Martin Plath, Andreas Svensson, Dustin Wilgers, Pam Willis, and Kyle Young for sending data, and David Shuker, Zen Lewis and Tom Price for helpful discussions.

## Supplementary methods

### LITERATURE SEARCHES

I performed literature searches using the online databases Web of Science & Scopus on the 29^th^ October 2018. All searches included the following search terms relating to the social and physical environment:

- “context dependent” OR “environment dependent” OR density OR “encounter rate” OR “sperm competition risk” OR audience OR “number of options” OR “number of choices” OR “sex ratio” OR predat* OR season OR ((cost* OR constraint*) AND (travel* OR time OR movement*)) NOT human

Additionally, searches included one of four keyword combinations relating to mating behaviour:

- (mate OR mating) AND (choice OR preference* OR choosiness OR rejection)
- courtship OR courting OR “sexual signalling”
- mate AND (sampl* OR search*)
- “species recognition” OR “mate recognition” OR “reproductive isolation” OR (conspecific* AND discriminat*) OR ((mate OR mating) AND (hybridisation OR reinforcement)) OR (mating AND (heterospecific* OR interspecific*))

I performed these four searches using both databases, on the same day.

### CRITERIA FOR STUDY INCLUSION

#### MATING BEHAVIOURS

I classed as sexual signalling any behaviours that the authors suggest function to attract a mate. This includes long range signalling when mates are not obviously nearby (such as calling by male crickets or anurans), and close-range courtship that functions to advertise mate quality and achieve mating. This means that measurements of signalling did not require that mates or sexual stimuli are encountered, whereas measurements of responsiveness and choosiness did. Importantly, if signalling behaviour was shown to be preferentially directed towards a specific mate or phenotype then it was instead classed as choosiness (see below). This category included estimates of display frequency, display duration, display rate, display latency, display intensity (e.g. calling amplitude), and proportion of individuals displaying.

I classed as responsiveness any mating behaviours (with the exception of courtship, see above) summed across all mate options or stimuli. I especially included passive attraction to mates or stimuli (e.g. when a mate option is not actively courting), or responses to mates in the early stages of courtship. When such behaviours could be shown to be directed towards a specific mate, or type of mate, they were instead classed as choosiness (see below). I classed measurements of mating or mating rate as either responsiveness or choosiness depending on several criteria. First, if mating frequency was linked directly to a mate phenotype it was classed as choosiness. Second, classification depended on the sex role of the focal sex- a behaviour was classed as choosiness if it was linked to the choosy sex, and responsiveness if it was linked to the courting sex. I excluded cases were it was difficult to attribute choice to either sex. One exception to this was any case where mating frequency was recorded after both sexes experienced the same change in the environment. This behaviour was classed as responsiveness, and assigned to its own sex category (‘both sexes’).

The choosiness category included any behavioural measure that could interpreted as reflecting the strength of a mating preference, including: mating outcomes, proxy behavioural measures, and sexual signalling when compared between different options. Preferences were often linked explicitly to a trait (either a specific stimuli or a mate phenotype), but this did not need to be the case- a subject that consistently prefers one mate option over another can be said to have a strong preference even if the exact traits being chosen are not recorded. I classed active acceptance/rejection behaviours of choosers in response to being courted, especially when they lead to mating, as choosiness rather than responsiveness. This category also included behaviours that could be said to reflect mate sampling, such as the number of mates visited or the number of times a subject switched between two options. This is because, all else being equal, high sampling will lead to the same outcome as high choosiness- a reduction in mating rate.

#### ENVIRONMENTAL FACTORS

I sorted environmental factors into seven categories: predation risk, population density, adult sex ratio, operational sex ratio, travel cost, time cost, and variation in mate quality.

##### Predation risk

This category consists of studies comparing prey behaviour across contexts which differ in the perceived risk of predation. I included studies which tested both direct and indirect risk factors. I classed predation risk as direct when prey were tested in the presence and absence of a live predator. I classed all other types of test as indirect, and this included studies comparing prey behaviour in relation to the level of predator chemical or acoustic cues, the presence of artificial (model) predators, and the light level during tests for acoustic preferences (which don’t influence signal detection but do influence the likelihood of predation by visually hunting predators). I also included studies examining the risk of parasitism by parasitoids.

##### Population density

This category consisted of studies comparing mating behaviour at different population densities, while controlling for the sex ratio perceived by subjects. In most cases, the sex ratio was equal at all densities. Perceived density differed due to variation in the number of conspecifics experienced, the size of experimental arenas, the spacing of mate options, or the time delay between conspecific encounters. In all cases, studies needed to examine per capita levels of behaviour, rather than the overall frequencies of behaviours, which is confounded by the number of individuals present at different densities. Importantly, I did not include cases in which population density could influence the amount of resources available to subjects, as this could potentially influence individual condition (Cotton *et al*., 2006).

##### Adult sex ratio

This category included any study comparing mating behaviour under conditions that varied the number of available mates. The sex ratio differed due to variation is the number of males and females in a group, the number of potential mates or rivals encountered, the number of rivals encountered, whether mates are encountered sequentially or simultaneously (Dougherty & Shuker, 2015), and the time delay between mate encounters. Note that for studies examining the expression of visual courtship signals, I only included studies in which at least one mate was present in all sex ratio treatments.

##### Operational sex ratio (OSR)

This category included studies comparing mating behaviour across contexts that alter the number of mating or breeding opportunities whilst keeping the actual number of opposite-sex individuals constant. Included studies considered the OSR to differ due to changes in the number of reproductively active conspecifics, the availability of nesting sites, and the resources available.

##### Travel cost

This category consisted of studies considering two factors. The first is the strength of water flow in fishes, which influences the energetic costs needed for swimming. The second is sound level for acoustic stimuli, which could influence how far away subjects perceive stimuli to be. For this latter factor, I did not include studies that measured choosiness or responsiveness using latency data, as latency is confounded by the distance needed to travel in this experimental setup. Studies comparing behaviour in relation to the distance between mates were considered as a density effect rather than a travel cost effect.

##### Time cost

This category includes studies that compare mating behaviour under conditions that could influence the time available for mating or successful breeding. This could be influenced by the day of the breeding season, the time in the reproductive cycle, or the generation in bivoltine insects. Studies comparing behaviour over the breeding season were assigned to this category unless other factors (such as OSR) were explicitly shown to vary over the season. I only included studies examining short-term time costs, rather than long-term changes associated with animal age, as this time cost may be confounded with other state-dependent effects when comparing individuals of different ages (Cotton et al., 2006).

##### Variation in mate quality

This category includes studies comparing mating behaviour between subjects that experienced differing degrees of mate quality variation. The attribution of mate quality was determined by the study authors, and was in relation to either body condition or age. Importantly, I excluded studies that did not control for the average mate quality experienced by subjects. This category only applies to choosiness and responsiveness.

### EFFECT SIZE EXTRACTION AND CODING

I used the correlation coefficient r as the measure of effect size. In this analysis, the effect size represents the difference or change in a behaviour due to the environment. Larger values therefore represent a greater difference in behaviour across different contexts. An effect size of zero indicates no difference in behaviour across contexts, but is blind to the magnitude across contexts. The vast majority of effect sizes (649 out of 697) were calculated by comparing behaviour expressed in two environmental contexts. When studies compared behaviours between two contexts, I first calculated the standardised mean difference, known as Hedges’ *d* (Hedges & Olkin, 1985; Koricheva et al., 2013), which was then converted to *r*. Hedges’ *d* was calculated either using reported means and standard deviations, or converted from common statistical tests (e.g. t tests, chi-squared, Mann-Whitney U test). I note that in most cases the traditional distinction when calculating the standardised mean difference between control and treatment groups did not apply. This is because it is not typically possible to assign one treatment as a ‘control’, given that there is no such thing as an unmanipulated or ‘standard’ environment. Instead, I compared behaviours between contexts, with the direction of comparison determined in relation to the environmental conditions (see main text for more details). When studies compared behaviours between more than three contexts, or across individuals rather than groups, I converted the results directly to the correlation coefficient *r*. When studies compared behaviours between more than two contexts, and where correlations were not appropriate, I performed multiple pairwise tests, or compared the two most extreme contexts.

A measure of choosiness requires that preferences be compared between two (or more) options. Therefore, to compare choosiness across contexts, I used two types of data. For frequency data, an effect size could be calculated from a 2 x 2 frequency table tallying the numbers of choices for one option over another in each context. For continuous data, preferences had to be presented as a single value, such as a difference score or some other metric, to facilitate comparison of multiple contexts. A difference score is the difference in preference between the preferred and non-preferred option.

I extracted effect sizes from both independent measures and repeated-measures experimental designs. In cases where effect sizes came from a repeated-measures design (for example using a test statistic from a paired t-test), I assumed a correlation of 0.5 between the two measures of behaviour. In this case, the effect size calculations are identical to those derived from independent-measures tests (Koricheva et al., 2013). A larger correlation between measures leads to a larger effect size estimate, and so I chose 0.5 in the absence of any reported correlation estimates, so as not to over-estimate these effect sizes. Note that in general the repeatability of mating preferences and courtship behaviour is generally low (Bell et al., 2009; Rosenthal, 2017).

### PHYLOGENY CREATION

I constructed a phylogenetic supertree for each of the six datasets primarily using the Open Tree of Life database (Hinchliff et al., 2015). However, several species were either missing from the OTL database or had an uncertain placement, resulting in polytomies in the tree. I resolved these issues manually by searching for published phylogenies that included the species if possible, or closely related species. When multiple trees were available I used the most recently published source. I successfully resolved all polytomies. I used Zhang et al. (2018) to establish the position of *Onthophagus binodis*. For the relationship among Araneae I used Wheeler et al. (2017). For the relationship among Lycosidae I used Piacentini & Ramirez (2019). For the relationship among *Shizocosa* species I used Stratton (2005) & Gilman et al. (2018). For the relationship among Tetigiionidae I used Mugleston et al. (2013). For the relationship among Gryllidae I used Chintauan-Marquier et al. (2016). For the relationship among *Gryllus* species I used Huang et al. (2000). For the relationship among Poeciliidae I used Reznick et al. (2017). For the relationship among *Poecilia* species I used Ptacek & Breden (1998) and Schories et al. (2009).

### HETEROGENEITY ANALYSIS

I used I^2^ as a measure of heterogeneity of effect sizes (Higgins et al., 2003). As the traditional formulation of I^2^ is unsuitable for multilevel meta-analyses, I used the extended method of Nakagawa & Santos (2012) to partition heterogeneity into its study-specific, species-specific and phylogenetic effects. To do this I used the MCMCglmm v2.29 R package (Hadfield, 2010). As in Metafor, I ran a multilevel random-effects model, with study, species, phylogeny and observation ID as random factors, on the three full (zeroes included) datasets. Each model was run for three million iterations, with a burn-in period of two million iterations and a thinning interval of 500. For all models I used an inverse-Wishart prior (V= 1, ν= 0.002).

## Supplementary tables and figures

**Figure S1.**
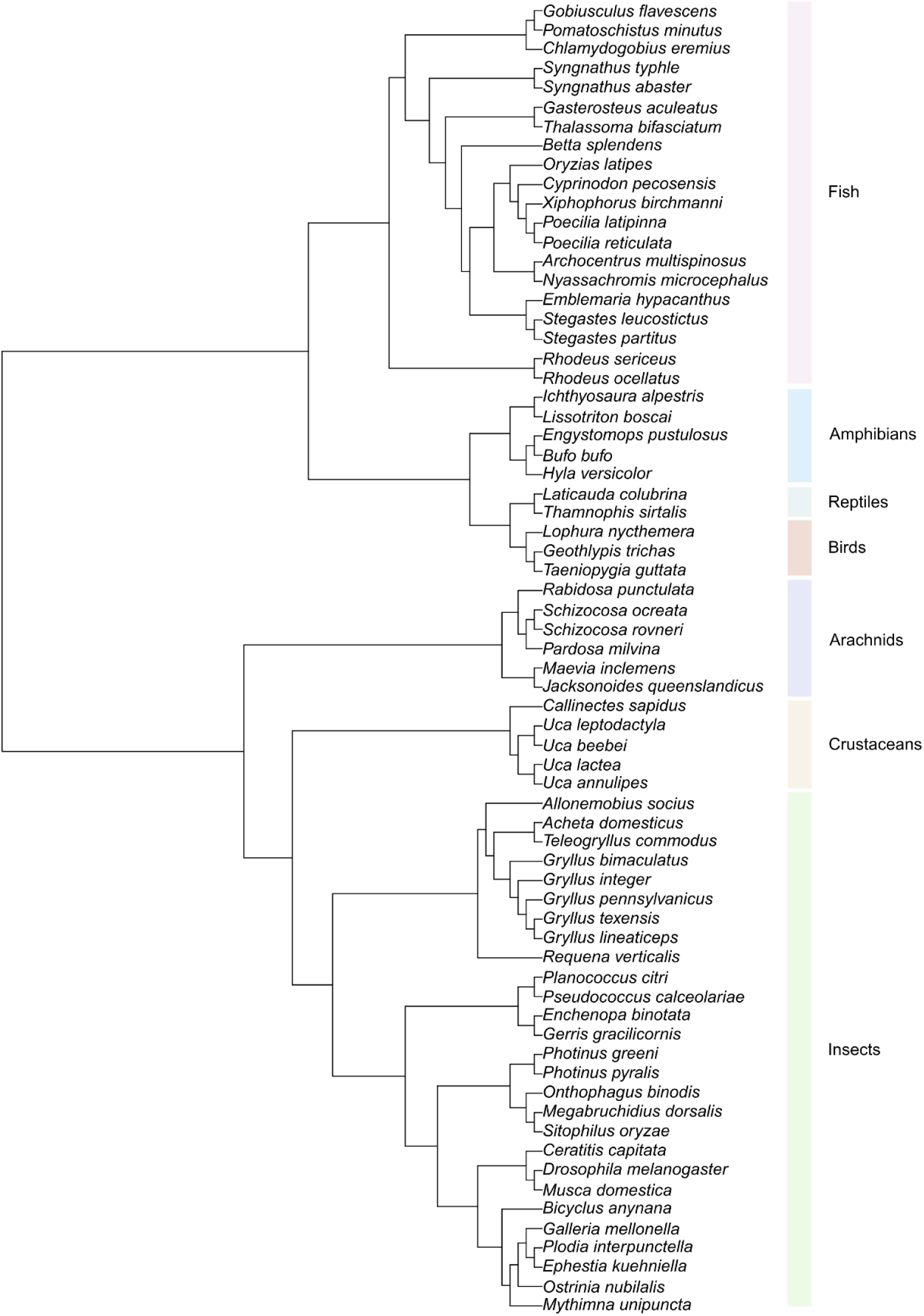
Phylogenetic relationships among the 68 species in the full sexual signalling dataset (zeroes included). Note that the branch lengths are not time-calibrated. Instead, all branch lengths were set to one and then standardised using Grafen’s method.

**Figure S2.**
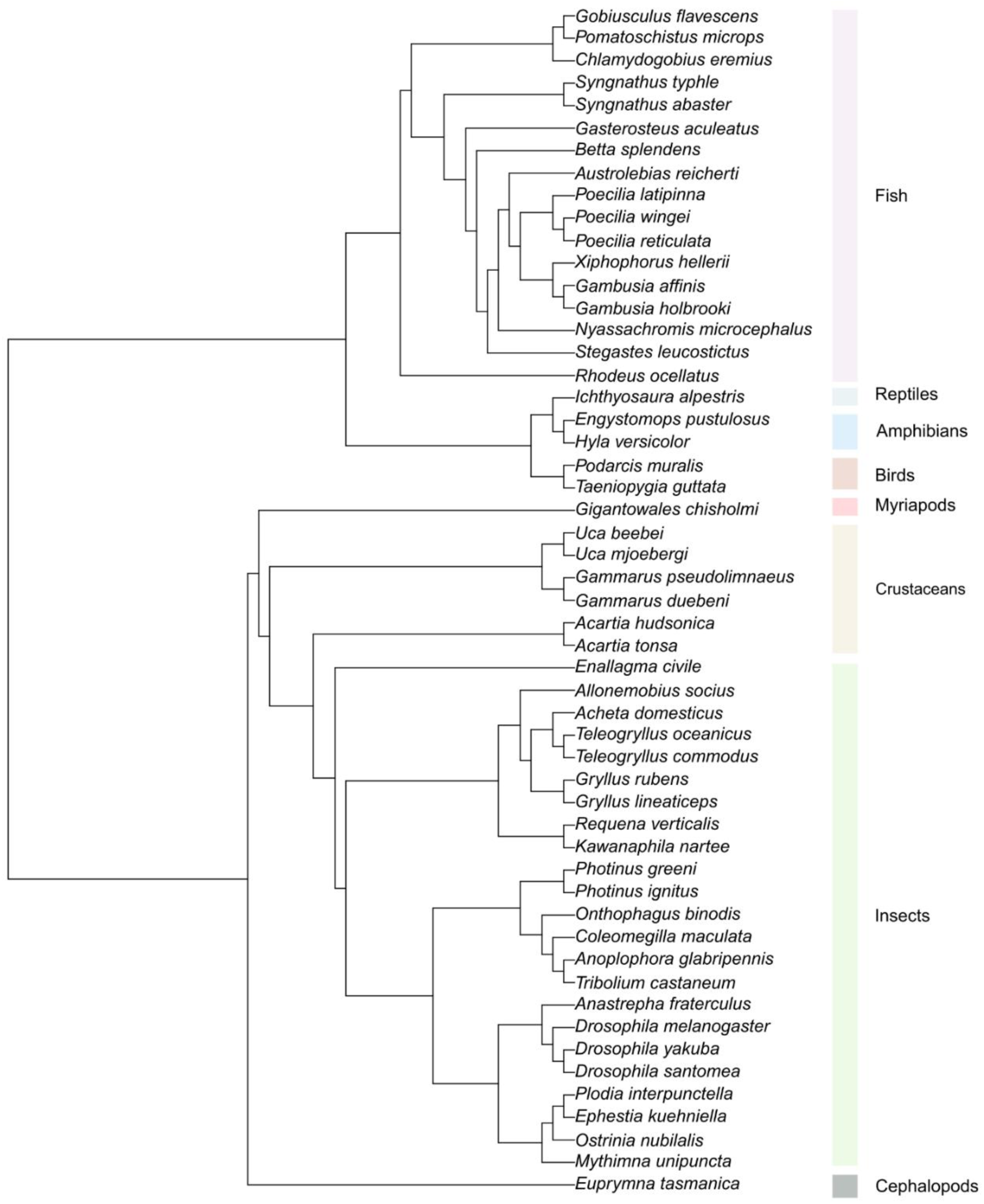
Phylogenetic relationships among the 53 species in the full responsiveness dataset (zeroes included). Note that the branch lengths are not time-calibrated. Instead, all branch lengths were set to one and then standardised using Grafen’s method.

**Figure S3.**
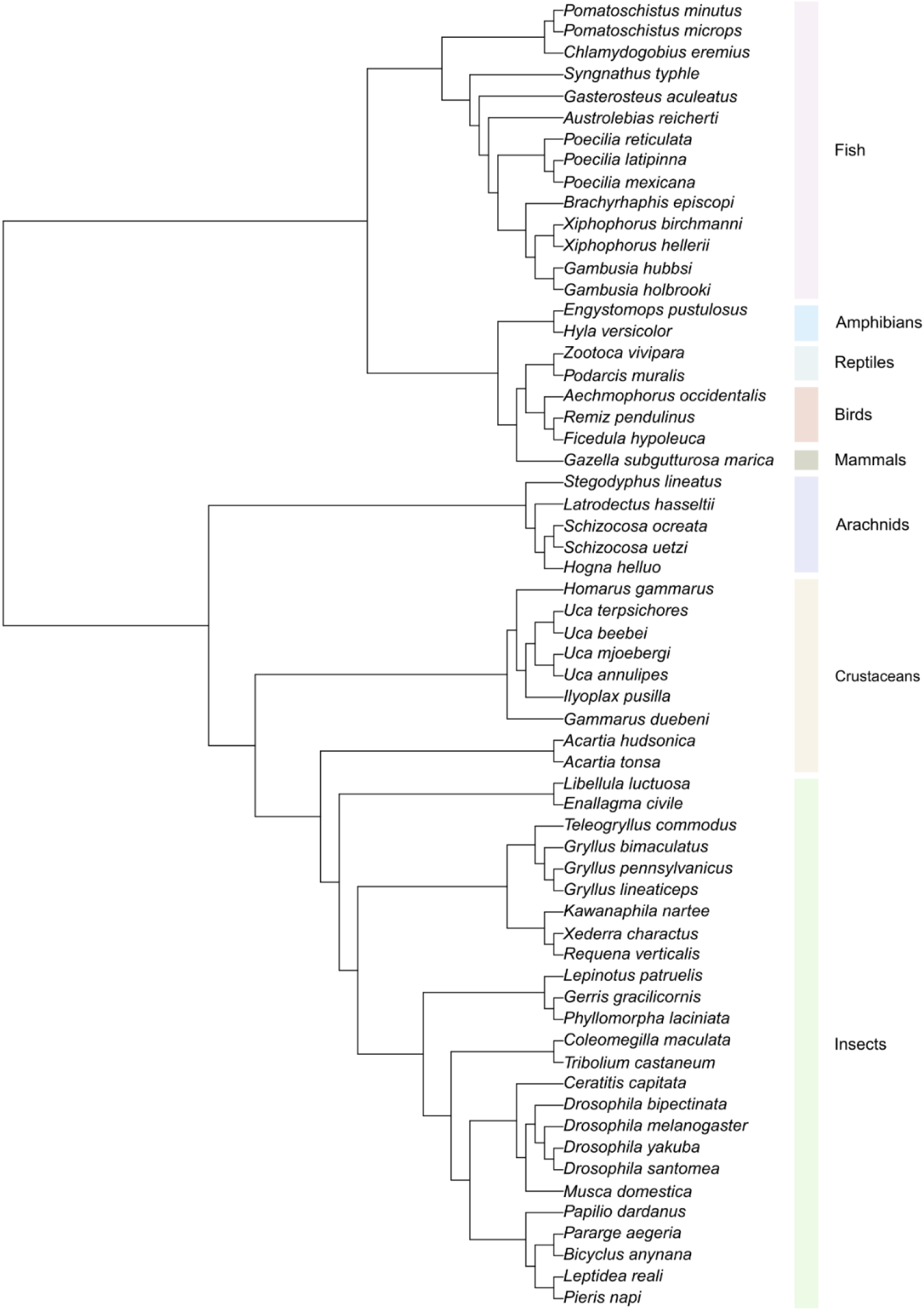
Phylogenetic relationships among the 61 species in the full choosiness dataset (zeroes included). Note that the branch lengths are not time-calibrated. Instead, all branch lengths were set to one and then standardised using Grafen’s method.

**Table S1.** Sample sizes and mean effect size estimates (correlation coefficient r, plus 95% confidence intervals) for each moderator category for the (full) sexual signalling dataset. Estimates were obtained using meta-regressions with four random factors and a single fixed factor.

| Moderator | Category | N Effect sizes | N Studies | N Species | Mean effect size ($r$) | 95% CI Lower | 95% CI Upper |
| --- | --- | --- | --- | --- | --- | --- | --- |
| Taxonomic class | Crustaceans | 11 | 7 | 5 | -0.276 | -0.735 | 0.357 |
|  | Arachnids | 33 | 9 | 6 | 0.156 | -0.437 | 0.655 |
|  | Insects | 72 | 31 | 27 | 0.050 | -0.450 | 0.527 |
|  | Fish | 113 | 53 | 20 | 0.082 | -0.440 | 0.563 |
|  | Amphibians | 17 | 7 | 5 | 0.202 | -0.424 | 0.698 |
|  | Reptiles | 7 | 3 | 2 | 0.213 | -0.462 | 0.732 |
|  | Birds | 7 | 4 | 3 | 0.249 | -0.434 | 0.751 |
| Focal sex | Male | 230 | 108 | 63 | 0.080 | -0.108 | 0.262 |
|  | Female | 24 | 10 | 11 | -0.017 | -0.266 | 0.235 |
|  | Both | 6 | 3 | 3 | 0.018 | -0.388 | 0.418 |
| Environmental factor | Density | 33 | 14 | 9 | 0.062 | -0.195 | 0.310 |
|  | Sex ratio | 136 | 69 | 47 | 0.031 | -0.170 | 0.229 |
|  | OSR | 3 | 2 | 2 | 0.103 | -0.419 | 0.573 |
|  | Predation | 87 | 37 | 25 | 0.141 | -0.075 | 0.344 |
|  | Travel cost | 1 | 1 | 1 | 0.164 | -0.576 | 0.756 |
| Factor timing | Before test | 54 | 29 | 19 | 0.101 | -0.093 | 0.287 |
|  | During test | 173 | 75 | 51 | 0.083 | -0.079 | 0.241 |
|  | Both | 33 | 13 | 9 | -0.085 | -0.317 | 0.157 |
| Factor variation | Manipulated | 235 | 100 | 60 | 0.058 | -0.177 | 0.286 |
|  | Natural | 25 | 17 | 13 | 0.159 | -0.128 | 0.421 |
| Animal origin | Lab-reared | 118 | 56 | 30 | 0.043 | -0.210 | 0.291 |
|  | Wild | 30 | 19 | 16 | 0.057 | -0.232 | 0.337 |
|  | Wild-caught | 112 | 39 | 28 | 0.099 | -0.153 | 0.339 |
| Signalling modality | Acoustic | 65 | 24 | 20 | 0.048 | -0.174 | 0.265 |
|  | Chemical | 2 | 2 | 2 | 0.327 | -0.367 | 0.787 |
|  | Mixed | 23 | 11 | 8 | 0.241 | -0.045 | 0.490 |
|  | Tactile | 8 | 4 | 3 | 0.107 | -0.284 | 0.467 |
|  | Visual | 162 | 73 | 37 | 0.043 | -0.141 | 0.224 |
| Signalling type | Short-range | 172 | 71 | 45 | 0.066 | -0.132 | 0.259 |
|  | Long-range | 88 | 46 | 30 | 0.080 | -0.131 | 0.285 |

**Table S2.** Sample sizes and mean effect size estimates (correlation coefficient r, plus 95% confidence intervals) for each moderator category for the (full) responsiveness dataset. Estimates were obtained using meta-regressions with four random factors and a single fixed factor.

| Moderator | Category | N Effect sizes | N Studies | N Species | Mean effect size ($r$) | 95% CI Lower | 95% CI Upper |
| --- | --- | --- | --- | --- | --- | --- | --- |
| Taxonomic class | Cephalopods | 1 | 1 | 1 | -0.135 | -0.801 | 0.680 |
|  | Myriapods | 1 | 1 | 1 | -0.530 | -0.891 | 0.243 |
|  | Crustaceans | 8 | 6 | 6 | -0.035 | -0.403 | 0.343 |
|  | Insects | 54 | 24 | 23 | 0.014 | -0.259 | 0.285 |
|  | Fish | 104 | 48 | 17 | -0.019 | -0.289 | 0.254 |
|  | Reptiles | 5 | 3 | 3 | 0.057 | -0.424 | 0.512 |
|  | Amphibians | 1 | 1 | 1 | 0.000 | -0.685 | 0.685 |
|  | Birds | 2 | 2 | 1 | 0.175 | -0.451 | 0.686 |
| Focal sex | Male | 78 | 45 | 27 | -0.013 | -0.118 | 0.092 |
|  | Female | 80 | 43 | 24 | 0.025 | -0.082 | 0.130 |
|  | Both | 18 | 9 | 11 | -0.082 | -0.308 | 0.153 |
| Environmental factor | Density | 24 | 8 | 9 | 0.122 | -0.106 | 0.338 |
|  | Sex ratio | 75 | 44 | 32 | -0.070 | -0.180 | 0.042 |
|  | OSR | 3 | 1 | 1 | 0.049 | -0.363 | 0.445 |
|  | Predation | 65 | 32 | 18 | 0.089 | -0.040 | 0.216 |
|  | Travel cost | 3 | 2 | 2 | -0.540 | -0.821 | -0.050 |
|  | Variation in mate quality | 6 | 2 | 2 | -0.113 | -0.458 | 0.261 |
| Factor timing | Before test | 67 | 37 | 26 | 0.046 | -0.067 | 0.158 |
|  | During test | 88 | 46 | 32 | 0.003 | -0.102 | 0.108 |
|  | Both | 21 | 10 | 8 | -0.199 | -0.411 | 0.033 |
| Factor variation | Manipulated | 153 | 74 | 46 | -0.002 | -0.086 | 0.083 |
|  | Natural | 23 | 13 | 10 | -0.011 | -0.197 | 0.176 |
| Animal origin | Lab-reared | 99 | 48 | 27 | 0.040 | -0.064 | 0.142 |
|  | Wild | 16 | 10 | 9 | 0.094 | -0.147 | 0.325 |
|  | Wild-caught | 61 | 28 | 21 | -0.112 | -0.246 | 0.027 |
| Preference measure | Matings | 19 | 11 | 12 | 0.039 | -0.161 | 0.236 |
|  | Proxy | 157 | 80 | 46 | -0.008 | -0.089 | 0.074 |
| Stimuli type | Mate quality | 161 | 83 | 53 | -0.002 | -0.081 | 0.078 |
|  | Species recognition | 15 | 5 | 7 | -0.029 | -0.232 | 0.176 |

**Table S3.** Sample sizes and mean effect size estimates (correlation coefficient r, plus 95% confidence intervals) for each moderator category for the (full) choosiness dataset. Estimates were obtained using meta-regressions with four random factors and a single fixed factor.

| Moderator | Category | N Effect sizes | N Studies | N Species | Mean effect size ($r$) | 95% CI Lower | 95% CI Upper |
| --- | --- | --- | --- | --- | --- | --- | --- |
| Taxonomic class | Crustaceans | 27 | 10 | 9 | 0.030 | -0.130 | 0.189 |
|  | Arachnids | 13 | 6 | 5 | 0.043 | -0.168 | 0.251 |
|  | Insects | 78 | 27 | 25 | 0.112 | 0.016 | 0.205 |
|  | Fish | 118 | 50 | 14 | 0.140 | 0.037 | 0.240 |
|  | Amphibians | 9 | 4 | 2 | 0.167 | -0.117 | 0.426 |
|  | Reptiles | 5 | 3 | 2 | -0.082 | -0.386 | 0.237 |
|  | Birds | 9 | 3 | 3 | 0.049 | -0.245 | 0.334 |
|  | Mammals | 2 | 1 | 1 | 0.000 | -0.521 | 0.521 |
| Focal sex | Male | 96 | 41 | 28 | 0.133 | 0.062 | 0.202 |
|  | Female | 159 | 72 | 48 | 0.080 | 0.021 | 0.139 |
|  | Both | 6 | 3 | 4 | 0.232 | 0.085 | 0.369 |
| Environmental factor | Density | 28 | 13 | 13 | 0.199 | 0.100 | 0.293 |
|  | Sex ratio | 122 | 52 | 39 | 0.073 | 0.002 | 0.143 |
|  | OSR | 9 | 5 | 5 | 0.178 | -0.020 | 0.363 |
|  | Predation | 64 | 29 | 17 | 0.122 | 0.019 | 0.222 |
|  | Travel cost | 10 | 4 | 3 | 0.123 | -0.147 | 0.375 |
|  | Time cost | 17 | 6 | 6 | 0.033 | -0.172 | 0.236 |
|  | Variation in mate quality | 11 | 2 | 2 | 0.031 | -0.232 | 0.290 |
| Factor timing | Before test | 95 | 43 | 29 | 0.082 | 0.003 | 0.160 |
|  | During test | 138 | 55 | 33 | 0.113 | 0.039 | 0.184 |
|  | Both | 25 | 9 | 8 | 0.108 | -0.051 | 0.263 |
| Factor variation | Manipulated | 209 | 84 | 47 | 0.100 | 0.039 | 0.159 |
|  | Natural | 52 | 22 | 19 | 0.095 | -0.011 | 0.198 |
| Animal origin | Lab-reared | 118 | 44 | 25 | 0.058 | -0.027 | 0.142 |
|  | Wild | 28 | 15 | 13 | 0.101 | -0.041 | 0.239 |
|  | Wild-caught | 112 | 45 | 26 | 0.134 | 0.053 | 0.212 |
|  | Mixed | 3 | 2 | 2 | 0.071 | -0.318 | 0.439 |
| Preference measure | Matings | 92 | 31 | 29 | 0.087 | 0.005 | 0.168 |
|  | Proxy | 169 | 83 | 46 | 0.104 | 0.041 | 0.167 |
| Stimuli type | Mate quality | 236 | 99 | 59 | 0.093 | 0.036 | 0.149 |
|  | Species recognition | 25 | 9 | 10 | 0.160 | 0.042 | 0.274 |

**Figure S4.**
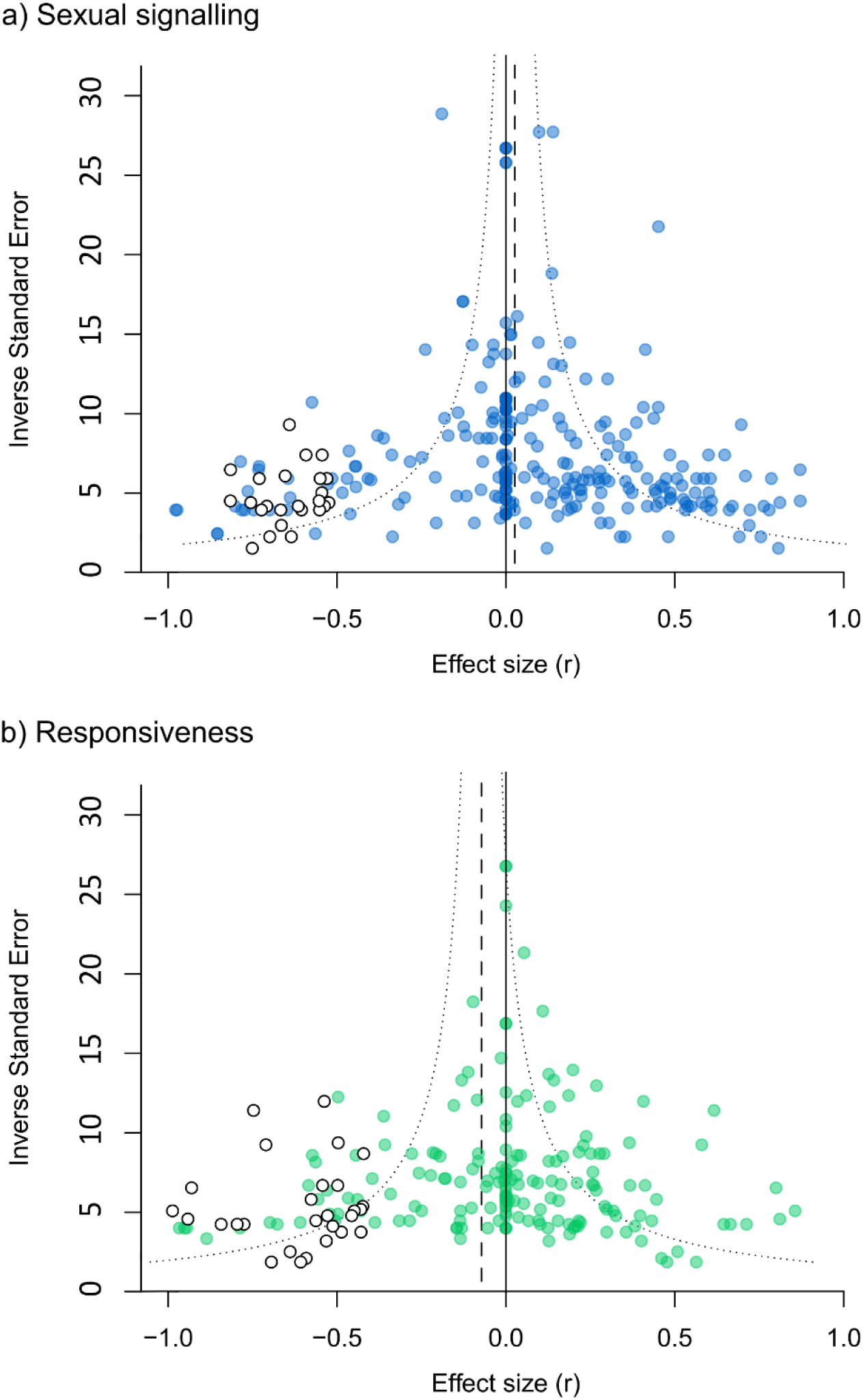
Trim-and-fill results for a) sexual signalling, and b) responsiveness. Filled circles show the observed effect sizes, and open circles show the imputed ‘missing’ effect sizes. The dotted line shows the mean effect size estimate after adding these ‘missing’ effect sizes.

**Figure S5.**
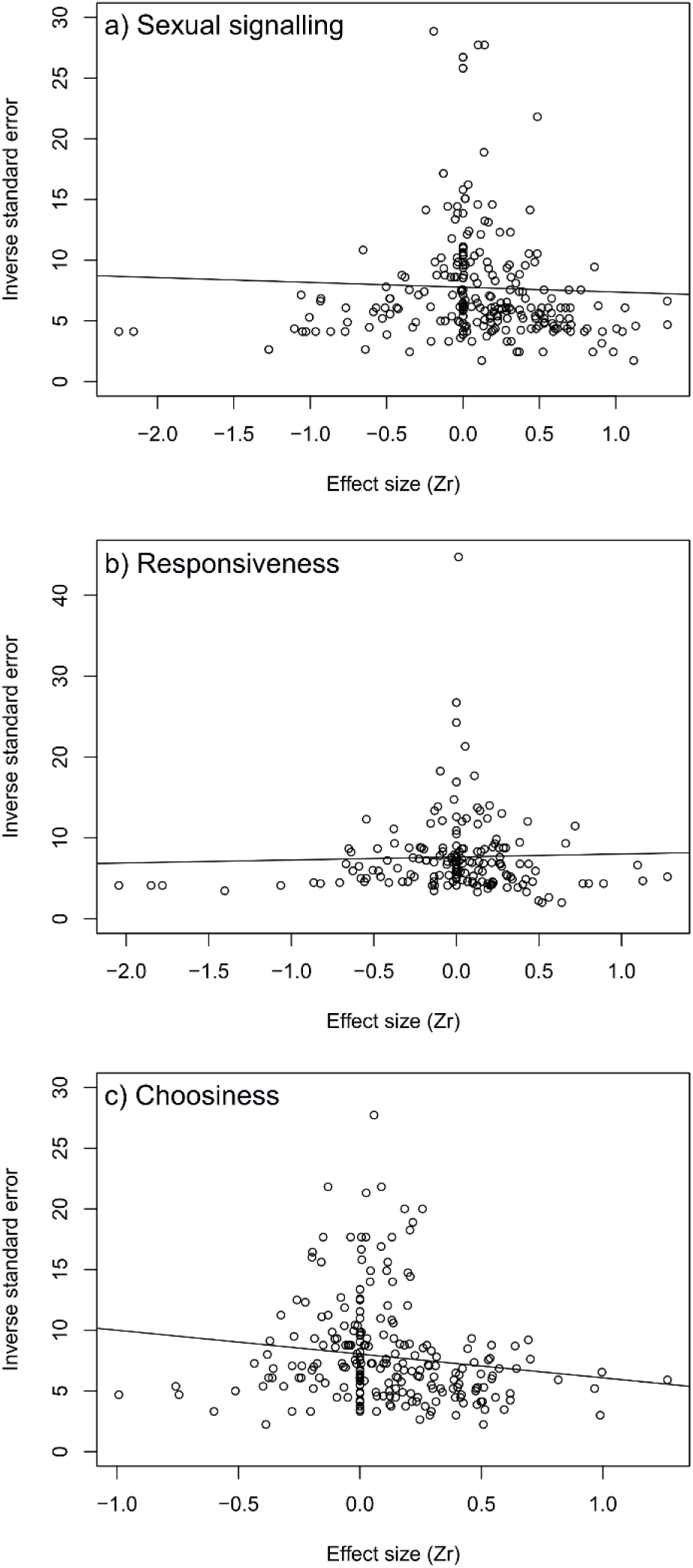
Scatter-plots (plus regression line) showing the relationship between effect size (*Zr*) and inverse standard error (a measure of study precision) for a) sexual signalling, b) responsiveness and c) choosiness.

